# Insurgence and worldwide diffusion of genomic variants in SARS-CoV-2 genomes

**DOI:** 10.1101/2020.04.30.071027

**Authors:** Francesco Comandatore, Alice Chiodi, Paolo Gabrieli, Gherard Batisti Biffignandi, Matteo Perini, Stefano Ricagno, Elia Mascolo, Greta Petazzoni, Matteo Ramazzotti, Sara Giordana Rimoldi, Maria Rita Gismondo, Valeria Micheli, Davide Sassera, Stefano Gaiarsa, Claudio Bandi, Matteo Brilli

**Affiliations:** Dipartimento di Scienze Biomediche e Cliniche L. Sacco, Università degli Studi di Milano, Milano, Italia; Centro Ricerca Pediatrica Romeo ed Enrica Invernizzi, Università degli Studi di Milano, Milano, Italia; Dipartimento di Bioscienze, Università degli Studi di Milano, Milano, Italia; Dipartimento di Scienze Clinico-Chirurgiche,Diagnostiche e Pediatriche, Università degli Studi di Pavia, Pavia, Italia; Dipartimento di Scienze Biomediche, Sperimentali e Cliniche, Università degli Studi di Firenze, Firenze; Laboratorio di Microbiologia, Virologia e Diagnostica della Bioemergenza, ASST Fatebenefratelli Sacco, Milano, Italia; Dipartimento di Biologia e Biotecnologie L. Spallanzani, Università degli Studi di Pavia, Pavia, Italia; Policlinico San Matteo Pavia Fondazione IRCCS, Unità di Microbiologia e Virologia, Pavia, Italia

## Abstract

The SARS-CoV-2 pandemic that we are currently experiencing is exerting a massive toll both in human lives and economic impact. One of the challenges we must face is to try to understand if and how different variants of the virus emerge and change their frequency in time. Such information can be extremely valuable as it may indicate shifts in aggressiveness, and it could provide useful information to trace the spread of the virus in the population. In this work we identified and traced over time 7 amino acid variants that are present with high frequency in Italy and Europe, but that were absent or present at very low frequencies during the first stages of the epidemic in China and the initial reports in Europe. The analysis of these variants helps defining 6 phylogenetic clades that are currently spreading throughout the world with changes in frequency that are sometimes very fast and dramatic. In the absence of conclusive data at the time of writing, we discuss whether the spread of the variants may be due to a prominent founder effect or if it indicates an adaptive advantage.

## Introduction

The worldwide fast spread of SARS-CoV-2 virus during the first months of this year has caused 316,169 deaths with more than 4,731,458 confirmed cases since the first reports of novel pneumonia (COVID-19) in Wuhan, Hubei province, China (Zhou et al. 2020; Wu et al. 2020) up to May 19 2020 (WHO 2020b). The virus belongs to the beta-coronavirus, and it is the seventh coronavirus known to infect humans, causing severe respiratory and systemic disorders (Rothan and Byrareddy 2020), with a basic R0 index estimated to range from 1.4 to over 6 (Lu et al. 2020). The closest known relatives of SARS-CoV-2 circulate in animals, specifically bats or pangolins (Zhang, Wu, and Zhang 2020), suggesting that an animal virus crossed species boundaries to efficiently infect humans, possibly through multiple passages in intermediate animal hosts, even though the transmission route has not been yet identified to date (Andersen et al. 2020).

Traces of the history of the spread are present in the viral genome and comparative genomics approaches can therefore be used to understand how viruses can adapt to multiple hosts, uncovering key signatures of this adaptation (Andersen et al. 2020; Wan et al. 2020), and to trace the infection routes of the virus. At the same time, genomic studies can help tracing viral variants that may be geographically restricted and/or may account for different levels of infectivity and mortality in humans. These variants might arise during the spread of the epidemic, as viruses are known for their high frequency of mutation, particularly in single stranded RNA viruses – as in the case of SARS-CoV-2 (Sanjuán and Domingo-Calap 2016), which has a single, positive-strand RNA genome.

Randomly generated variants can then spread in the population, due to stochastic reasons (i.e. founder effect, drift) or as a consequence of positive selection exerted by intrinsic biological features (such as the level of virus infectivity and its transmission rate), or extrinsic factors such as use of antivirals or reactions by the immune system or other defence mechanisms put in place by the host (Di Giorgio et al. 2020). Therefore, haplotype(s) present at the beginning of the epidemic spread can change in time; sometimes novel variants can overcome ancestral ones, and this can be a consequence of different levels of aggressiveness but also of mechanisms beyond selection. If the different variants are identical in terms of their ability to infect and replicate in the host, country-specific switches in haplotype frequency with respect to the most common haplotypes can depend on the very first haplotype(s) arriving in the country (Provine 2004). Instead, when haplotype frequency changes globally, then the hypothesis of differential aggressiveness becomes more probable, indicating that the novel variants may be better adapted to infect human hosts; however, even in this case, the complexity of global human mobility or sampling bias, may originate patterns that may seem causal but are not.

Here, we present a comprehensive study of the coding sequences from SARS-CoV-2 genome sequences isolated since the beginning of the epidemic. Italy was the first European country registering non-imported COVID-19 cases requiring hospitalization. The first registered case of COVID-19 was on February 20, 2020, a young man in the Lombardy region in Northern Italy, diagnosed with atypical pneumonia (Livingston and Bucher 2020). In the next 24 hours 36 more cases were detected, and as of April 27th 2020, Italy registered 197,675 cases with 26,644 deaths, one of the highest toll in Europe and in the world. WHO classifies the spreading occurring in Italy as community transmission, indicating that the country is experiencing large outbreaks of local transmission with no possibility to trace transmission chains between the cases, and with multiple and unrelated clusters of transmission. The high mortality rate observed in Italy, especially at the beginning of the epidemic raised the question whether Italian strain(s) might have increased aggressiveness. While the increased mortality observed in Italy could be explained by the fact that, at the beginning of the epidemic, swabs were performed only on individuals showing up at a hospital, distorting the sampling toward a group of symptomatic individuals devoid of healthy carriers, we cannot exclude that Italian haplotype frequency are partially different, at least genetically. To have a better insight on the history and spread of the COVID-19 pandemic in Italy and thanks to the sequences deposited in the Gisaid database, we identified 7 non synonymous mutations that are differentially frequent in Italian SARS-CoV-2 strains respect to strains circulating globally. These mutations are enriched in Italy, but present in strains from other countries as well, as shown by tracing their relative frequency in time both globally and in different countries, therefore we traced their distribution worldwide and complemented with a phylogenetic analysis to understand how the variants are related.

## Material and Methods

### Genome sequences and extraction of coding sequences

Genomes were downloaded from the Gisaid.org repository on April, 10, 2020, and a second time on April, 28, 2020, and are listed, together with reference to the submitting laboratories, in Supplementary Table 1. We extracted coding sequences using a strategy based on tblastn (Camacho et al. 2009) comparisons using as queries all the proteins from the SARS-CoV-2 reference sequence deposited in NCBI (Accession MN908947). After the comparison, the coordinates of the blast alignments on each genome were used to extract the coding sequence. Nucleotide sequences were then aligned and translated, and the alignments were manually checked for the presence of frameshifts, and manually edited. Alignments were manually edited to remove partial and poorly aligned sequences, resulting in a variable number of sequences per alignment. This manual curation resulted in alignments containing a minimum number of 2262 sequences for ORF3a and a maximum of 5585 sequences for the Nucleocapsid protein, with an average of 4222 sequences per alignment. We are aware that by removing sequences with gaps we are likely removing part of the genuine variability present (i.e. indels), however, we observed indels with such a low frequency that we attributed them mostly to sequencing/assembly errors and decided that the clear advantage of a stronger signal outweighs the possible disadvantages deriving from information loss. Additional analysis with high quality genome sequences will be necessary to evaluate whether indels represent an important source of variation. Regardless, this does not change the results of our analysis as the target of this work are point mutations.

### Identification of variable and differentially evolving sites in coding sequences

Sequences from the alignments were used to build amino acid frequency profiles by using the R-package seqinr (Charif and Jr 2007). Basically, for each protein encoded in the SARS-CoV-2 we obtained two profiles, one for Italian strains and one for the entire set of sequences. As the positions in the alignments for the two groups under examination are congruent, we can then calculate:

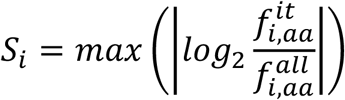

that is the maximum log ratio of the frequency (*f*) of all amino acids at a certain position *i* in the alignment in Italian (^it^) and total (^all^) sequences. Then we identify positions with S>0.5 that corresponds to the identification of positions where there is a frequency change in one of the residues of at least 2X. Variability in multi-alignments was quantified by calculating entropy 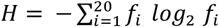 at each position of each manually curated protein multi-alignment. Mutual Information was calculated based on (Buck and Atchley 2005) to assess whether variants tend to co-occur frequently.

### Time profiles

To build updated time profiles of the identified variants, sequences were downloaded a second time from Gisaid.org on April, 28, 2020 (9075 sequences, only high quality and high coverage). Sampling dates were retrieved from the database and can be found in the Gisaid Sequence Acknowledgment table (Supplementary Table 1). Sequences for which the geographic origin and/or the data were not correctly defined (Supplementary Table 1), were removed, to obtain a database of about 8500 sequences, with some per-site variability due to the variable presence of Ns in the sequences. We calculated two different time series, one for the entire set of sequences available up to the moment of sequence download, and one by considering only sequences taken from predefined groups of countries (hereinafter the time series by country). In the latter, we merged nearby countries to increase the precision of the estimation of the relative frequency of variants within pre-defined time intervals. The total time range since the first sequence available (133 days) was split into 13 (9 for the analysis by country, as the number of sequences per interval is smaller) non-overlapping intervals of about 10 days (∼13 days for the time series by country). Interval duration was selected looking for a compromise among the number of sequences available on average per interval in the different groups and the time resolution of the time series. For each interval we calculated the frequency of the observed variants and the Shannon diversity index considering all concatenated coding sequences (not limited to the positions considered in this work). In Supplementary Table 2 we show the number of sequences used for each country for each interval.

### Phylogenetic analysis

We performed a phylogenetic analysis by concatenating nucleotide aligned on the basis of the manually curated amino acid alignments. We selected the best phylogenetic model for our alignment using ModelTest-NG (Darriba et al. 2020) (GTR+I+Г model) and then we performed Maximum Likelihood phylogenetic estimation in RaxML8 (Stamatakis 2014). The obtained phylogenetic tree was then visualized and integrated with additional information about variants and geographical origin using FigTree (https://github.com/rambaut/figtree/).

## Results and discussion

### Amino acid variants

Our analysis allowed us to identify 7 positions in four proteins that present drastic changes in amino acid frequencies when comparing Italian sequences with worldwide sequences available on Gisaid.org on April, 10, 2020 (Figure 1). However, we discovered that residues found at these positions are not peculiar of Italy and, as a matter of fact, they are the most variable genomic positions across available SARS-CoV-2 sequences (Supplementary Figure 1). Therefore, we decided to proceed with a worldwide analysis. The sites that we identified are:

**Figure 1.**
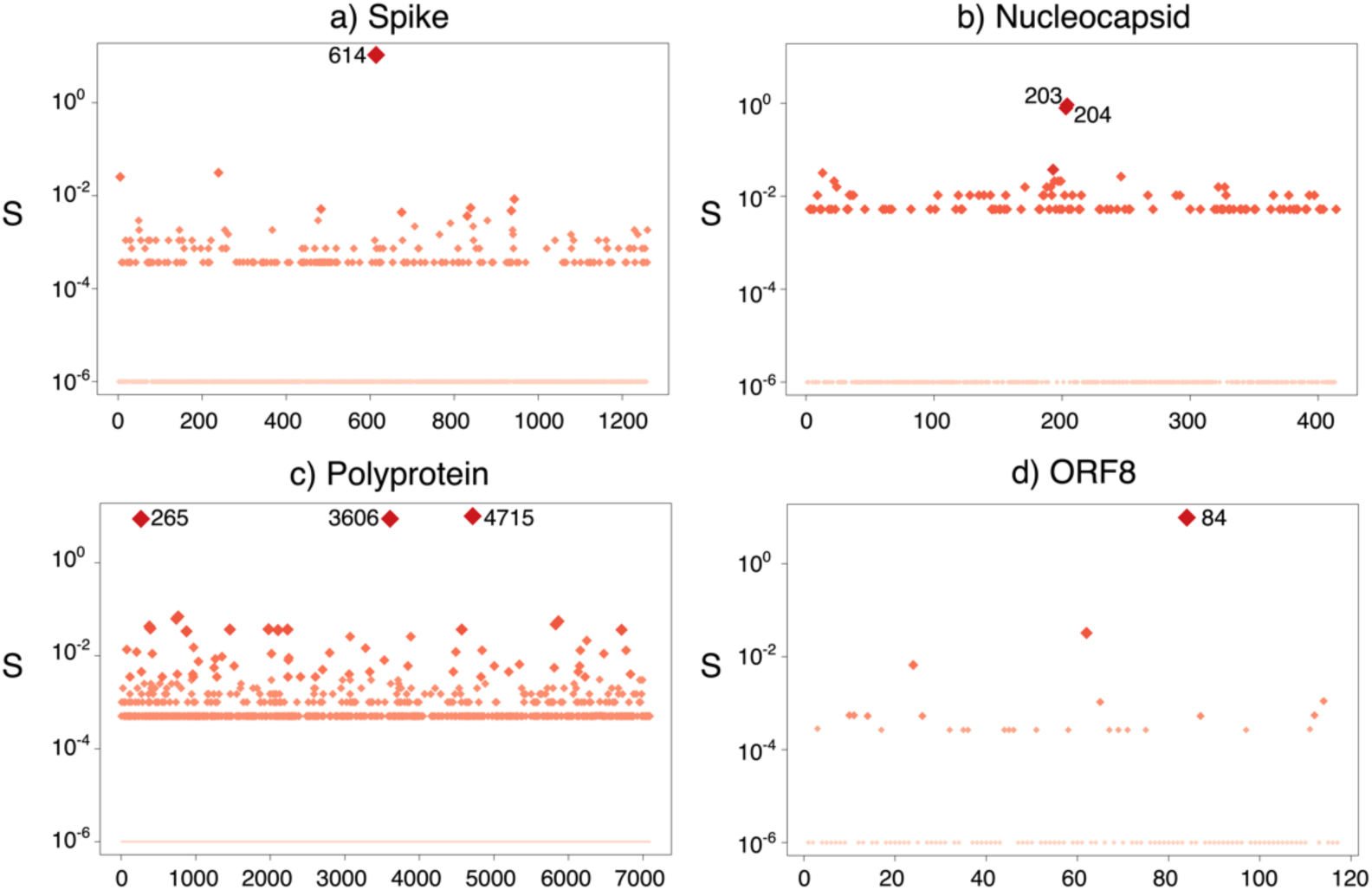
Positions identified as maximally different when contrasting amino acid frequencies in italian Vs the entire dataset. See Supplementary Figure 1 for a confirmation of the variability of these positions in the entire set of sequences. Note that to cope with zeros when taking the logarithm of the ratio of amino acid frequencies, we add a small number to every frequency that depends on the number of sequences in the two groups (italian and world). This makes so that positions with identical amino acid frequencies in the two groups do not get an S exactly equal to zero.

1. Position 614 in the Spike protein, where an Aspartate residue is found in high frequency in sequences obtained at the beginning of the pandemic. A variant, with a Glycine residue at the same position is however now very common in Europe, Italy in particular.
2. Positions 203 and 204 in the Nucleocapsid protein.
3. Positions 265, 3606 and 4715 in the Polyprotein.
4. Position 84 in Orf8.

During the preparation of this manuscript, we realized that some of the variants are also the topic of other papers and preprints predating this work e.g. (Korber et al. 2020; Banerjee et al. 2020; Somasundaram, Mondal, and Lawarde 2020; Begum et al. 2020; Bai et al. 2020; Yang et al. 2020; Pachetti et al. 2020). The fact that we identify some of the previously known mutations by using a different approach indicates that they represent a genuine signal rather than artefacts; however, this does not necessarily mean that their increase in time (see below) is a consequence of a correspondingly increased transmissibility rate or aggressiveness in general – for which at the moment there is no conclusive data as stated in most of the available preprints. In the following, we summarize a thorough bibliographical analysis looking for functional information associated with these positions or the domains they belong to. However, we stress once more that virus spreading is characterized by haplotype shifts that are not necessarily related to the performances of the haplotypes themselves – for instance genetic drift or founder effect – therefore without targeted functional studies on the variants it is difficult to ascertain whether one or the other explanation is more probable.

### Spike variant

D614 is located in the SD1 domain of the spike protein. It is positioned in a loop right after a beta-strand and close to a completely solvent exposed disordered region outside and downstream the ACE2 interaction domain (Wrapp et al. 2020; Walls et al. 2020). The resolution of the three available structures is not high enough to thoroughly discuss H-bond network. It is likely that D614 side chain does not establish close interactions with neighboring residues. Conformationally, D614 lies in the region of left-handed helices. Thus, from the structural point of view, the mutation D614G is likely to be neutral (no particularly strong interactions lost from the missing carboxylate group). On the other hand, the mutation might even be beneficial for overall protein stability: given the increased conformational freedom of Glycine, it is indeed a perfect residue in turns following beta structures. Other authors suggest a positive effect of D614G on the virus efficiency in binding the ACE2 receptor. Korber and colleagues (unpublished, (Korber et al. 2020)) suggest the possibility of structural changes in the protein or even an improved antibody-dependent enhancement effect.

### Nucleocapsid variants

Positions 203 and 204 belong to the linker region between the N- and C-terminal RNA binding domains. In particular they are positioned in the SR region named after the marked abundance in S and R residues (McBride, van Zyl, and Fielding 2014). This region plays a role in cell signaling and RNA binding. It is a thought to be a highly flexible and intrinsically disordered region (IDP) of the protein. While the mutation R203K is a conservative mutation in terms of charge and size of the residue side chain, G204R introduces a bulky charged residue in place of a G residue. However, even though the loss of a G may reduce the backbone flexibility, R residues are common in IDP regions and R204 might introduce a new charge which is compatible with the RNA binding properties of this domain. As said before, we feel that the change from RG to KR may indicate a requirement for an Arginine at that position, with a path RG to KG, that could be disadvantageous, to KR and back to a situation resembling the original haplotype.

### Polyprotein variants

In Coronaviruses, ORF1 is translated to yield the polyprotein that is processed by proteolysis with the production of intermediate and mature nonstructural proteins (nsps). In Figure 2 we highlight the position of the variant sites on the Polyprotein with respect to known domains, as defined by the Conserved Domain Database (Lu et al. 2020).

**Figure 2.**
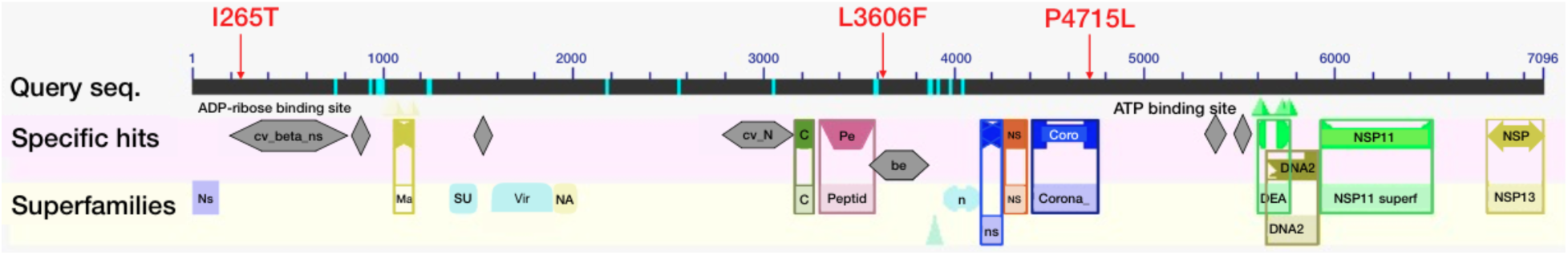
Position of the variant sites on the Polyprotein sequence, with indicated the domains as defined on the basis of the Conserved Domain Database (S. Lu et al. 2020).

Position 265 corresponds to residue 85 of the non-structural protein 2 (Nsp2) which is characterized by four transmembrane helices (TH) (Angeletti et al. 2020), whose role is still unknown. More in detail, position 85 lies at the N-terminal, which protrudes from the external face of the membrane. Interaction analysis of the corresponding protein from SARS virus shows that Nsp2 can form dimeric or multimeric complexes that can also involve additional viral proteins (Nsp3, Nsp4, Nsp6, Nsp8, Nsp11, Nsp16, ORF3a) (von Brunn et al. 2007) suggesting that Nsp2 might be involved in the viral life cycle. SARS Nsp2 also interact with host proteins, such as prohibitin 1 and 2 (Cornillez-Ty et al. 2009), that are involved in cell cycle progression, migration, differentiation, apoptosis and mitochondrial biogenesis (Fusaro et al. 2003; Merkwirth and Langer 2009; Rajalingam et al. 2005; Sun et al. 2004) suggesting it might be important to manipulate prominent host functions. Murine Hepatitis Virus and SARS-CoV strains deleted in this portion of the polyprotein (Graham et al. 2005) have a strongly reduced viral growth and RNA synthesis, but are not affected at the level of protein processing. Ectopic expression of the nucleotide sequence coding for Nsp2 in murine cells infected by strains missing Nsp2, allowed to detect its recruitment in viral complexes. It is therefore possible that Nsp2 is involved in global RNA synthesis, interaction with host proteins and pathogenesis (von Brunn et al. 2007). In infected cells, Nsp2 seems to be present in small vesicular foci, but in absence of additional viral proteins it localizes in cytoplasmic or nuclear membranes, without a specific target (Prentice et al. 2004; von Brunn et al. 2007). Unfortunately, mutagenesis experiments are still not available for SARS-CoV2, and therefore we still do not know whether the variant observed at this site has any effect on *in vivo* virus properties.

Position 3606 belongs to Nsp6, a protein that induces the formation of autophagosomes, which is a sign of starvation in uninfected cells (Cottam, Whelband, and Wileman 2014). Autophagosomes can act as an innate defense against viral infection but they can be hijacked and support the assembly of coronavirus replicase proteins (Orvedahl et al. 2007; Suhy, Giddings, and Kirkegaard 2000; Wileman 2006). Together with Nsp3 and Nsp4, Nsp6 moreover promotes the formation of the double membrane vesicles typically observed in SARS disease (Angelini et al. 2013). Nsp6 is a protein with 7 transmembrane domains (Oostra et al. 2008) and position 3606 lays in the luminal loop between the first and second hydrophobic transmembrane domains. In the wild type SARS-CoV-2 sequence, we find 3 Phenylalanine residues just before L3606. Instead, some of the variants in this paper have a stretch of 4 Phenylalanine, keeping this region highly hydrophobic.

Position 4715 belongs instead to the N-terminal domain of the RNA-directed RNA Polymerase (RdRp, also known as Nsp12). By inspecting its three-dimensional structure, SARS-CoV-2 RdRp displays a N-terminal nidovirus-like RdRp-associated nucleotidyltransferase domain (NiRAN) followed by an interface domain and then by the canonical palm and finger domain structure (Gao et al. 2020). P4715 (P323 according to Nsp12 numbering) is located in the alpha-beta interface domain which bridges NiRAN with the finger domain. Specifically, P4715 sits on a solvent exposed loop region in the groove formed between NiRAN and the finger domain. The mutation P4715L mutation does not change the non-polar nature of the side chain and from the conformational point of view the substitution from P to L should not result in specific structural adjustment of the loop. Thus, despite no information concerning mutations at the specific sites identified in this work, the three variant sites in the polyprotein belong to important functional regions of the sequence.

### Orf8 variant

Orf8 codes for a secreted accessory protein not directly involved in viral replication (Dediego et al. 2008; Yount et al. 2005; Tan et al., 2020). Homologs have been found in some beta-coronavirus, named Orf7a (Tan et al., 2020). Orf7a of SARS codes for a protein with a structure similar to immunoglobulin superfamily proteins, specifically the metazoan Ig involved in adhesion (Nelson et al. 2005; Hänel et al. 2006). The function of this protein is not entirely clear, but could be related to the modulation of the immune system. Studies have demonstrated that SARS-Orf7a protein localizes mainly in the perinuclear region of the host cell, where it interacts with BST-2 (bone marrow stromal antigen 2) preventing the glycosylation of BST-2 needed for functioning (Taylor et al. 2015). BST-2 inhibits the release of the virus, as observed in HIV-1 (Taylor et al. 2015) likely at the level of the endoplasmic reticulum-Golgi where it binds budding virions. Orf8 has a structure composed of a beta-sandwich fold with seven beta-stands, which highly resembles Orf7a’s structure, but it lacks the C-terminal transmembrane region and has an additional long insert between strands 3 and 4 which is supposed to be involved in peptide binding (Tan et al., 2020). Orf8 is a fast-evolving gene which, together with the absence of the transmembrane region and the presence of the insertion may suggest some functional divergence with respect to its ancestor (Tan et al., 2020). It has been proposed that Orf8 may have acquired a function similar to the adenoviral CR1 protein, which interferes with MHC molecules to attenuate the antigen presentation and therefore the capability of the host immune system to detect the virus. In this context, we notice that L84 lays within the Orf8 insert, indicating it may represent a further adaptation. Because direct functional information on this site is lacking at the moment, it is not possible to ascribe a functional adaptation to these variants; however, variability at this site might counteract MHC interference function, as a fast-evolving region in the insert may have been selected positively to facilitate the interaction with a fast-evolving host molecule (Tan et al., 2020).

### Time series of the variable sites

After having identified variant sites, we explored the time profile of the seven variable sites across all sequences (Figure 3) evidencing haplotype frequency changes for all of them between February and March, with five changes being moderate (Nucleocapsid R203G and G204R, Orf8 L84S, Polyprotein T265I and L3606F) and two more drastic, now representing the most common haplotypes of non-chinese recent sequences (Spike D614G and Polyprotein P4715L). The Spike variant, in particular, was sequenced only few times in China since the beginning of the epidemy, the first time in Zhejiang on January 24; However, in this country it never reached a significant frequency. Conversely, after the first sequencing of this variant in Germany on January 28 this variant started at very low-frequency and then became the most common haplotype at this position outside China. A similar situation is true for the haplotype with a Leucine in position 4715 of the Polyprotein – that rapidly increases since the beginning of March.

**Figure 3.**
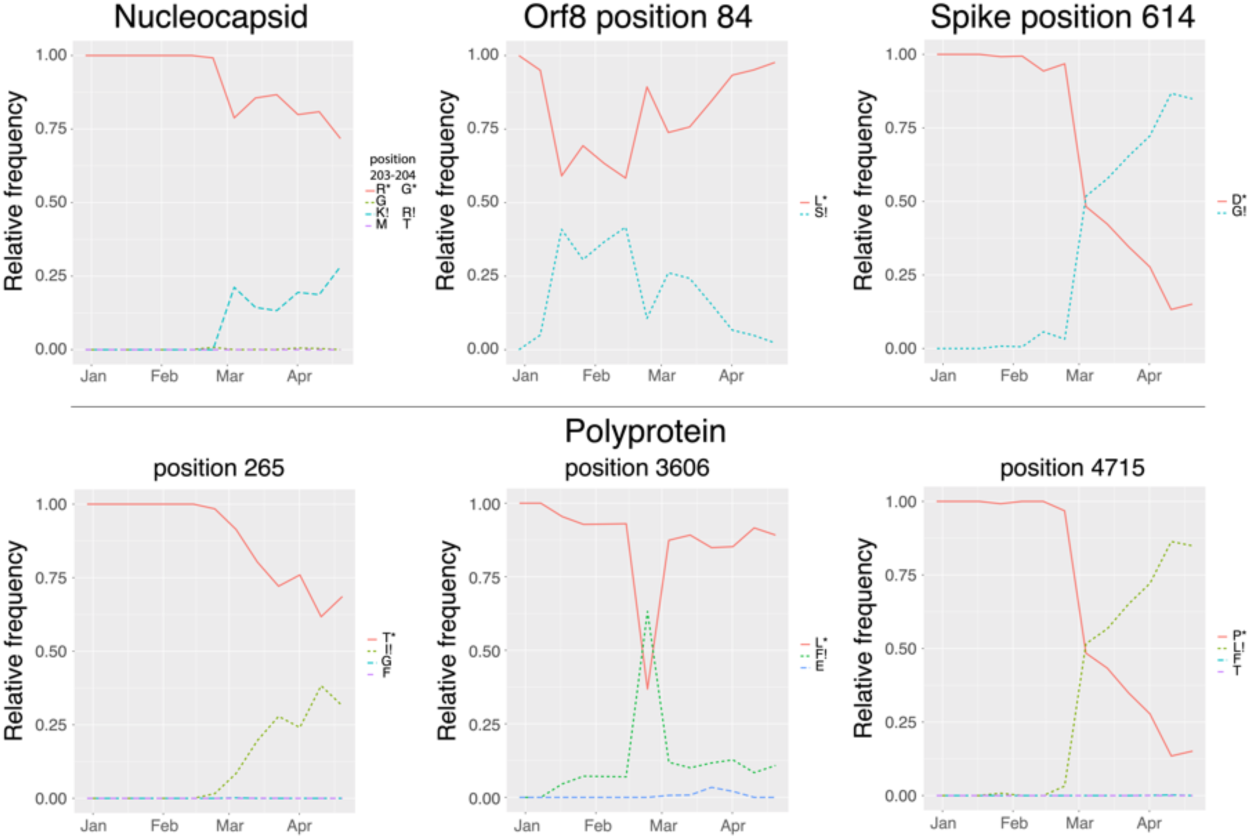
Time series for each variable site using sequences downloaded on April 28, 2020 (last sequence: April, 20). In the legend, we indicate the reference residue with an asterisk, and the variant on which we are focusing with an exclamation mark. Additional variants were identified in the sequence dataset downloaded on April, 28, but they never reach significant percentages, at least at the moment.

These patterns seem indicative of a functional role for at least some of the variants that undergo an increase in frequency, but in the absence of any functional test or experimental data, we cannot rule out that the observed frequency changes are a consequence of a founder effect in Europe followed by a spreading wave from Europe to countries where the epidemic started later. A founder effect, however, implies that the founder arrives first, while this is not always the case, at least for some of the variants and part of the countries. At the same time some data both reviewed and unreviewed start to be available, suggesting that D614G on the Spike might provide some advantage to the virus. Bai (Bai et al. 2020) and Brufsky (Brufsky 2020) observed a correlation of G614 with increased mortality, while Korber and coworkers (Korber et al. 2020) found a correlation between the presence of the mutation and a higher viral load in patients.

Position 614 on the Spike and position 4715 on the Polyprotein covary in a significant way (data not shown). The most likely explanation for this is the rapid sequential fixation of both mutations in the same strain, together with the absence of recombinations in between the two. The two adjacent sites in the Nucleocapsid sequences show a perfect agreement in the time profiles, strongly suggesting that they happened together or within a short time. These two mutations respectively remove and add an Arginine (passing from RG to KR); by considering the overall frequencies of the possible amino acid pairs at these positions, we suggest that the second Arginine may complement the loss of the first one. Indeed, RR is never observed, KG is extremely rare (relative frequency, r.f.= 0.00036), while most genomes either present the original pair RG (r.f.= 0.87464) or KR (r.f.= 0.12500). We speculate that this could indeed be a consequence of the non-neutrality of configurations with no Arginine at both positions, an information still not giving hints on the fitness of the fixed variant. Next,we reasoned that grouping all the sequences uploaded from different countries may provide a picture averaged over variable and more complicated situations that might characterize the evolution of the virus within different countries or geographical areas. We therefore explored the time profiles of the variants in different geographical macro-regions (Figure 4 and Figure 5) highlighting a highly heterogeneous situation. We also provide a movie illustrating the changes taking place in variant frequencies across overlapping time windows in macro-regions (Supplementary Movie 1), which provides a dynamical view of the time profiles of the variants.

**Figure 4.**
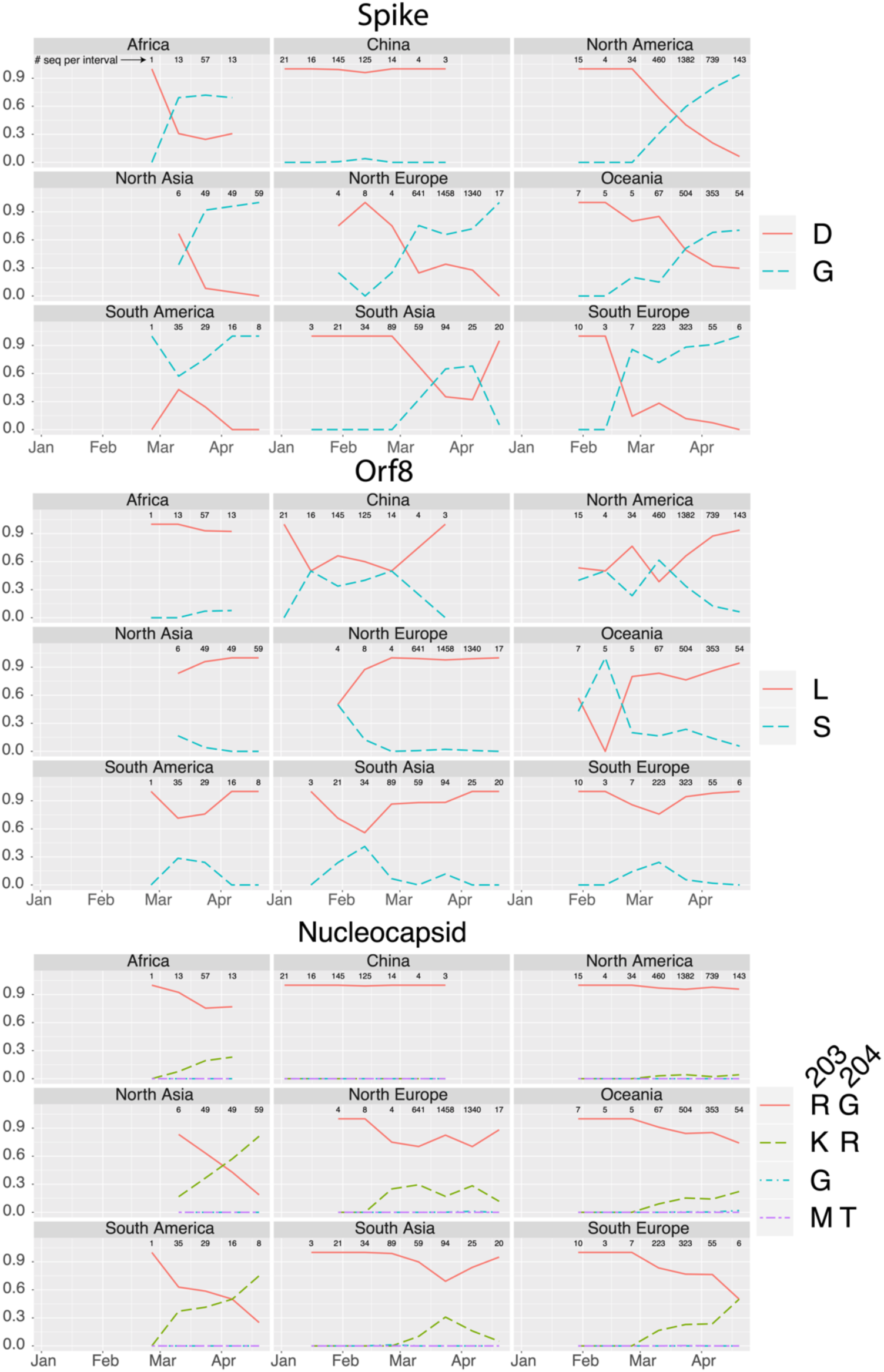
Time profiles the frequencies of single residue variants in geographic macro-regions for Spike, Nucleocapsid and Orf8. Please note that as the frequency profiles for variants in the Nucleocapsid are identical, we combined the two.

**Figure 5.**
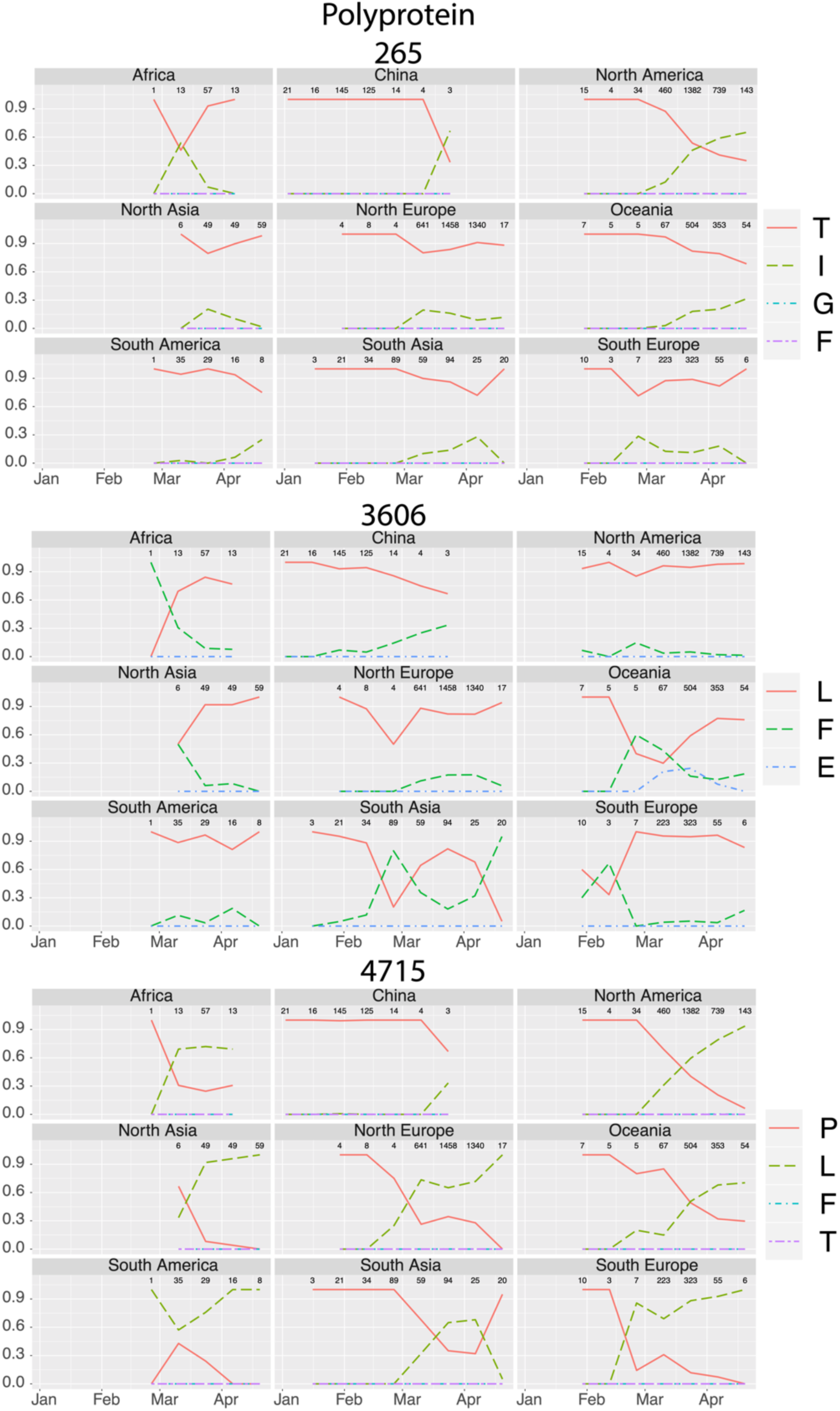
Time profiles of the frequencies of single residue variants in geographic macro-regions for the Polyprotein.

Figure 4 shows that D614G in China never reaches significant frequency while it increased quite rapidly in several areas where it arrived and where it often started at very low frequency with respect to the original haplotype. If a functional role for this mutation will be demonstrated, this pattern seems to indicate that different variants might have different fitness when interacting with different host’s haplotypes, i.e. in case Asian and European have different haplotypes concerning some of the proteins interacting with the Spike, like for instance Furin. To conclude, it is clear that since the first appearance of D614G and other of the above variants, their relative frequency underwent a significant increase in several countries, most of the time overcoming in prevalence the original variant(s), except for China, South Asia, South America and Africa. In Figure 6 we summarize the most recent situation, by using sequences from the interval April 10 to 20, 2020.

**Figure 6.**
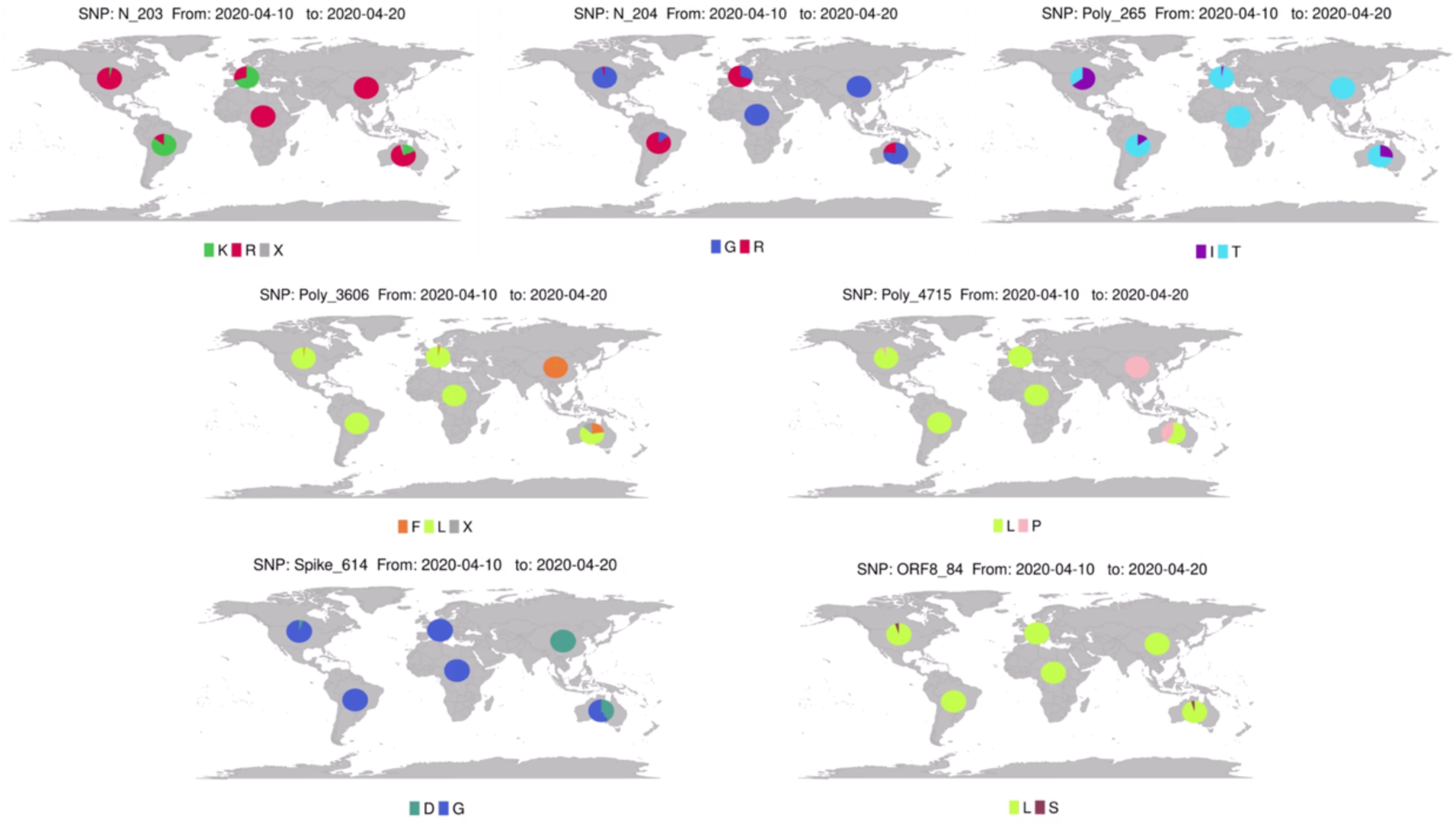
Abundance of the seven variants in the world, obtained with sequences available on Gisaid.org on April, 28, 2020, and relative to the interval April, 10-20; see also Supplementary Movie 1.

### Phylogenetic analysis and integration with variant information

The best phylogenetic model selected according to AIC and BIC was GTR+I+G, for which the estimated substitution rate matrix contains the following rates, relative to the G<->T rate, taken as unity: A<->C=0.34, A<->G=0.77, A<->T=0.32, **C<->T=100**. The extremely high rate for C to T (or better U, considering we are dealing with an RNA virus) is in agreement with the involvement of host APOBEC-like editing mechanisms, as proposed in recent works (Di Giorgio et al. 2020). Besides not being the focus of this paper, these rates provide further evidence that host’s mutagen systems may play an important role in the evolution of SARS-CoV-2 and its detection by the immune system. The phylogenetic tree integrated with additional information is reported in Figure 7, with different coloring schemes and together with an evolutionary model summarizing how the different variants configurations are likely related to each other. When considering the information about the residues found at the seven positions on which we are focusing (hereinafter variants configurations, VCs), as in Figure 7a, we find that the phylogenetic clades correspond to the six most frequent ones (accounting for over 96.3% of the sequences), that is RGTLPDL, RGTLPDS, RGTFPDL, KRTLLGL, RGTLLGL, RGILLGL – obtained by linking the variant residues in the order: protein Nucleocapsid sites 203, and 204, Polyprotein sites 265, 3606 and 4715, Spike site 614 and orf8 site 84; for this reason hereinafter we will use Clades A to F interchangeably with the VCs written above. We are aware that the correlation is not perfect, as can be seen by the presence of additional but low frequency variants within each clade, or the presence of VC misplaced with respect to their major clade. Misplaced sequences can be a consequence of insufficient phylogenetic signal or of convergent evolution. For instance, as anticipated above one of the Chinese sequences carrying Spike variant 614G (VC: RGTLPGS), is indeed contained within Clade B (RGTLPDS), indicating convergent evolution with the Spike D614G variant in Clades D, E, F, or maybe an artifact. However, the limited number of similar cases supports our simplification and the correspondence among clades in the tree and VCs.

**Figure 7.**
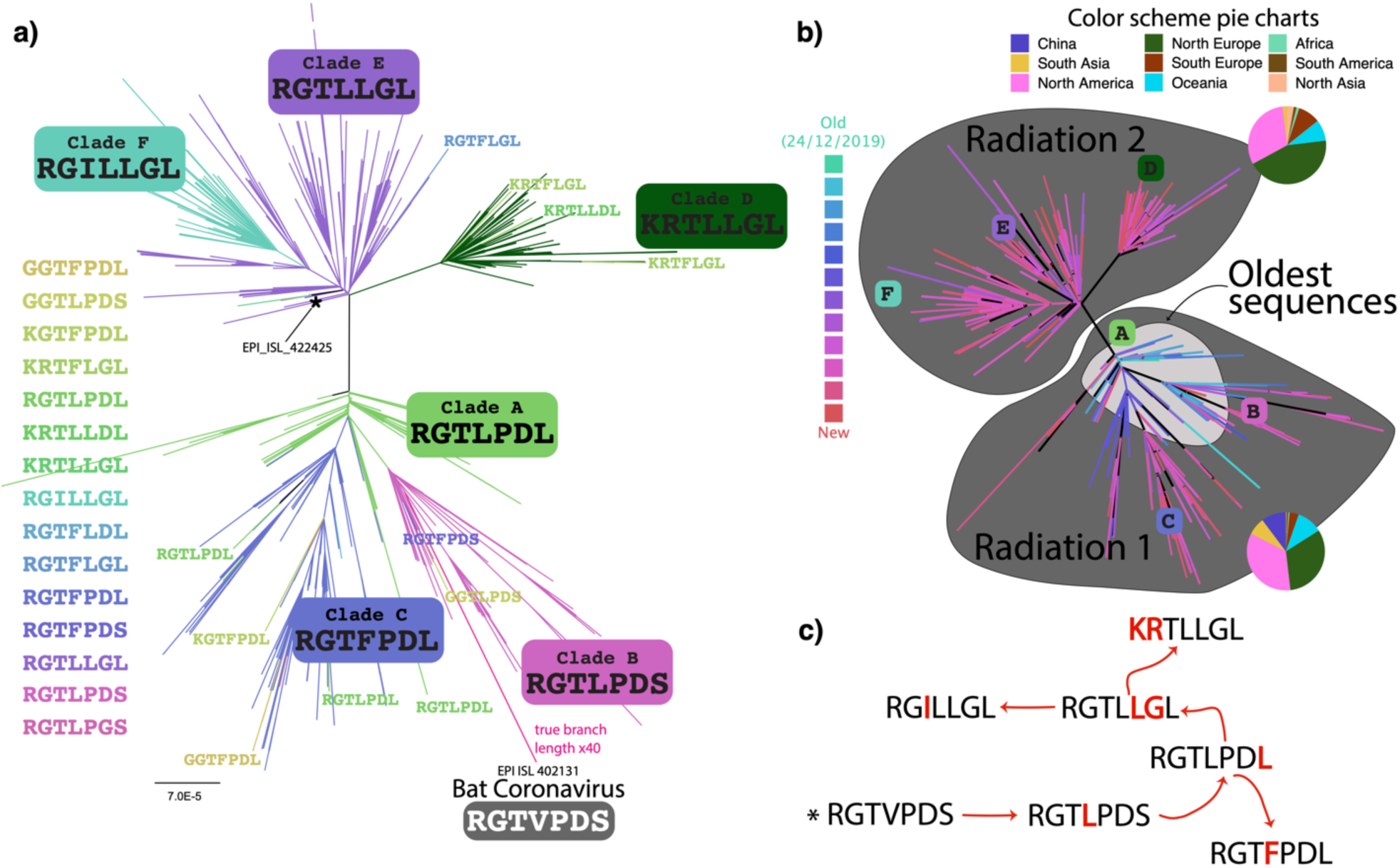
Phylogenetic analysis and integration with variants configurations. a) Phylogenetic tree with branches colored on the basis of tip VCs/Clades, showing the existence of 6 major “clades” corresponding to the six most abundant configurations of variants. b) The same phylogenetic tree, but with branches colored on the basis of the Collection Date of sequences. c) Inferred evolutionary model for variants configurations appearance.

Our multi-alignment contains one sequence annotated as Bat coronavirus (EPI_ISL_402131), which provides a rooting of the phylogenetic tree that falls in Clade B and indeed contains most of the sequences obtained in China at the beginning of the epidemy (Figure 6, panel b). The tree has two main “radiations”, one corresponding to variants configurations RGTLPDL, RGTLPDS, RGTFPDL and one to KRTLLGL, RGTLLGL, RGILLGL, therefore the main partition of the tree corresponds to the identity of residues at position 614 of the Spike and 4715 of the Polyprotein. The two radiations are present at different frequencies in the same countries, with the notable exception of chinese sequences within Radiation 2, except for EPI_ISL_422425, the only sequence with a RGTLLGL pattern sequenced in China that is also present in the phylogenetic tree (indicated in Figure 7a). This suggests that while most of the diversification of Radiation 1 took place in China and was then exported outside, the ancestor of Radiation 2 travelled outside China early to start a diversification in the countries where it arrived. This hypothesis is in agreement with the fact that EPI_ISL_422425 is placed very close to the root of the branch leading to Radiation 2, indicating it may indeed represent a strain closely related to the true Radiation 2 ancestor. These observations raise important questions (1) about the identity and movements of the first individual carrying the ancestral Radiation 2 variant outside China (2) on the timing of the epidemic, but most importantly (3) the reason why it did not increase in frequency in China but elsewhere.

Following the topology of the tree we propose an evolutionary model (Figure 7c) whereby the ancestral Bat variants configuration (RGTVPDS or any other present in the presently unknown animal host) evolved into RGTLPDS and from this to RGTLPDL. We stress that this likely took place through unobserved states/hosts as the root branch length is close to the total length of the tree and therefore we do not consider mutation V3606L in the Polyprotein full sequence as an adaptation to the human host. The latter originated RGTFPDL on one side, and the ancestor of Radiation 2 (RGTLLGL) that originated both KRTLLGL (through KG unfit strains?) and later RGILLGL.

Given the almost perfect agreement of the tree with the variants configurations, we analysed the geographical distribution and the time profiles of the seven sites at once, similar to what is done for MLST-based classification of pathogens. In Figure 8a we show the time profiles of the relative abundance of sequences belonging to the 6 clades, clearly showing not only the appearance and increase of the novel clades, but most importantly the gradual disappearance of sequences belonging to Clade A (see also Supplementary Movies 2 and 3). When focusing on single Clades across all macro-regions previously defined, we find a heterogeneous situation with different variants increasing in time in different countries. However, we can see that all VCs with a Proline at position 4715 of the Polyprotein and an Aspartate at position 614 of the Spike almost disappear in time, remaining abundant only in South Asia and Oceania; in the rest of the world, only VCs with a Leucine and a Glycine at those positions remain in the last interval of our time range, after replacing the existing VCs. This is indeed clear from Figure 8c showing that VC for Clade A (RGTLPDL) is no more present in the interval April 10 to 20, 2020. South Asia is particular because it is the only area where VCs of Radiation 2 (RGTLLGL,KRTLLGL,RGILLGL) are present for some time but then disappear with a re-increase of Clade C (RGTFPDL). This may suggest these variants have equal phenotypic characteristics, but as written above, we cannot exclude differential fitness depending on host’s haplotypes.

**Figure 8.**
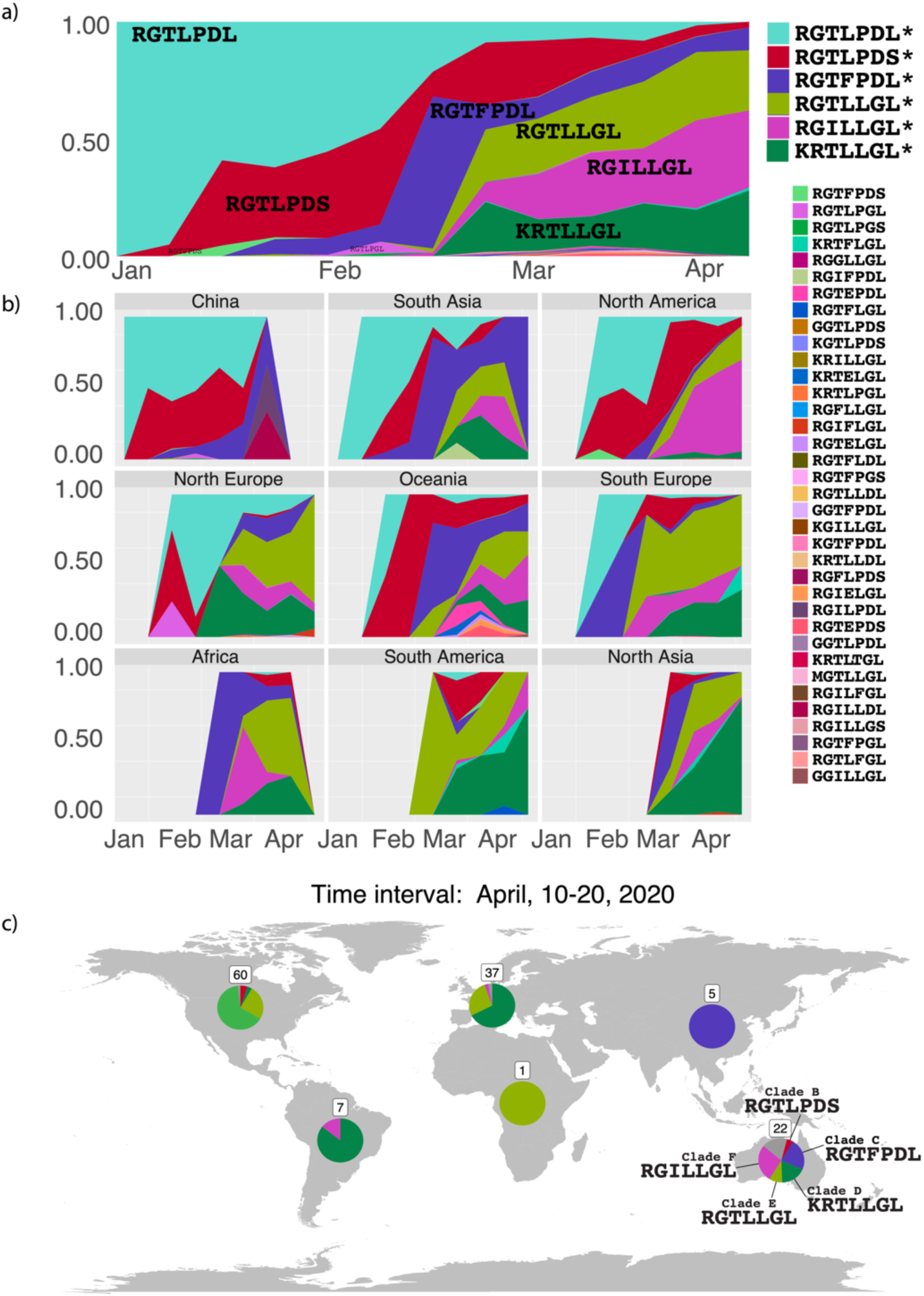
a) Time profile of sequences belonging to different clades/VCs considering all sequences together. y-axis correspond to the relative frequency in each time interval. Only the most frequent Clades are indicated; b) as above but for the different macro-regions. c) The relative abundances of sequences belonging to the different clades in different geographical macro-regions using sequences from the interval April 10-20, 2020. See also Supplementary Figure 2.

**Figure 9.**
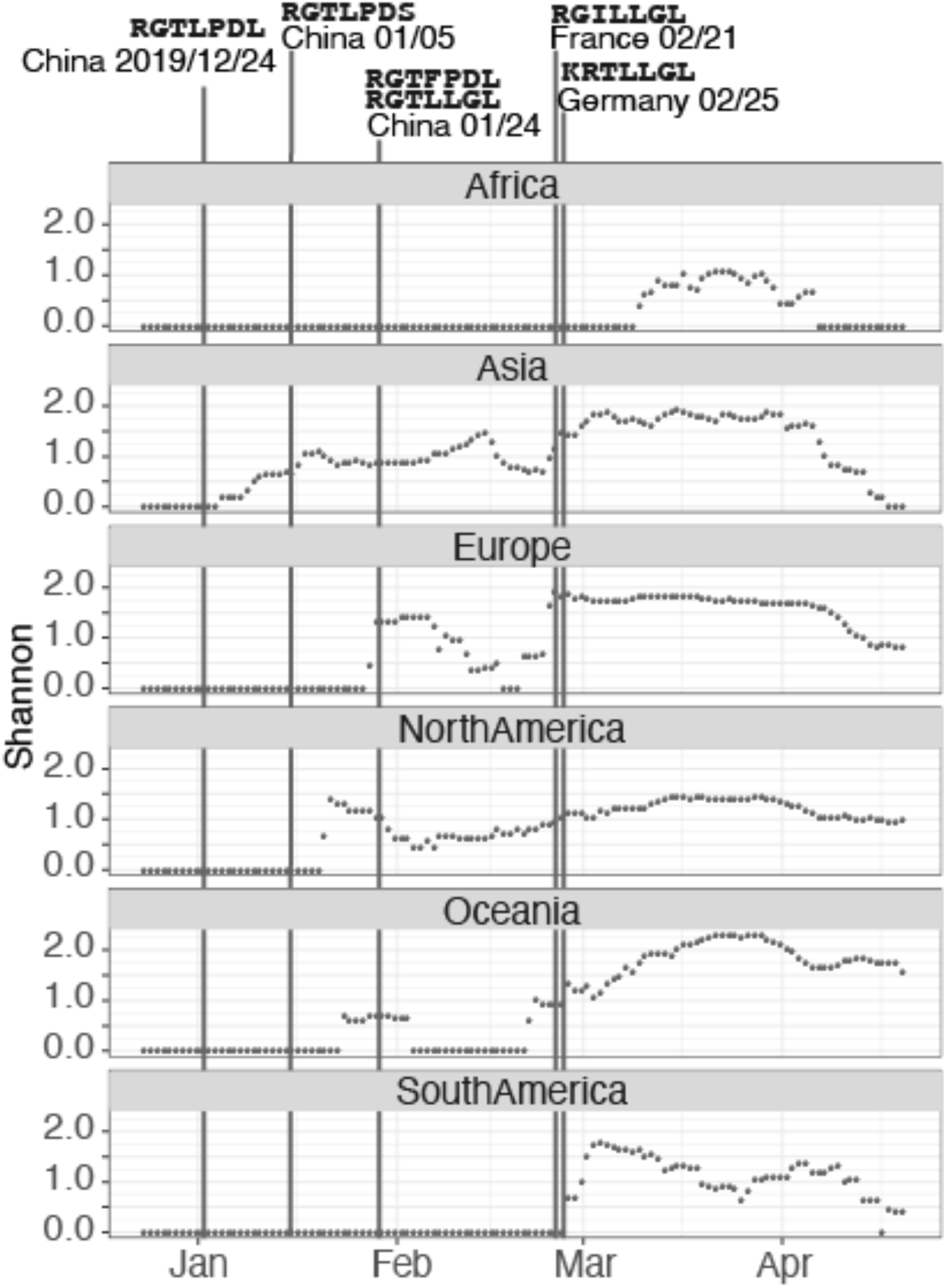
Shannon index as a function of time in different macro-regions, calculated for sequences within each time interval. We also indicate the first sampled sequence for each of the six clades. Variability increases when a new variant appears, then it stays more or less stable when the existing variants maintain their relative frequencies. However, if one of the variants becomes dominant, then variability decreases again after a peak. Increasing variability in time means that the arrival of novel variants continues steadily, or that the existing ones become more homogeneous in frequency.

By calculating the Shannon index for sequences within the same intervals of times used to track the changes in frequency of the variants, we were able to trace the evolution of diversity in different places. Peaks indicate the emergence (from outside or by evolution of pre-existing strains) of variants and their increase in frequency. Values maintaining a high Shannon index correspond to situations where different variants coexist at comparable frequencies, while a decrease after a peak means that after the appearance of one or more new variants / clades, one of them (novel or old) takes over – reducing the variability.

## Conclusion

In this work, we identified seven positions in coding sequences of the SARS-CoV-2 genome that are characterized by a different pattern of amino acids when comparing Italian sequences with the global trends around the world. Further analysis revealed that these sites are not peculiar of Italian strains, and that different combinations of these variants are present at varying relative frequencies in different geographic areas. We found that the combination of these residues identifies six abundant configurations that corresponds to 6 phylogenetic Clades and that cover over 96% of all sequences. This suggests that the characterization of these positions can represent a fast and portable method for the SARS-CoV-2 typing, but novel variants are emerging that might eventually take over the old ones through mutations at additional sites. Using this approach we were able to follow the evolution of the virus over time among continents, showing that the different clades evolved in different moments and that their frequencies vary among continents.

These sites are also the most variable among all available SARS-CoV-2 sequences, raising intriguing questions about their functional effects. Variants with a Leucine at position 4715 of the polyprotein together with a Glycine at position 614 of the Spike, underwent an increase in frequency since the end of January in most countries, overcoming the original haplotypes. Mutations that might affect the structure of the Spike protein are of primary interest, since many vaccine candidates and serological tests rely on the conformation of this protein (WHO 2020a). This and other works also explore the hypothesis that the variants may indeed provide a selective advantage to the virus. Clade prevalence in different countries could be used to check for mortality rate differences and association with variants, but as the rates depend on many other factors (different screening strategies, different ways to define an individual infected and so on), we feel premature discussing such correlations. Once the numbers will be standardized for different countries, this kind of associations, if present, will clearly emerge. Moreover, to really clarify these issues, experimental data is required, such as for instance in the form of Tn-insertion mutagenesis, as performed on other viruses in the past (Fulton et al. 2017) followed by competition experiments in *in vitro* cultures or the design of genomes carrying well-defined changes. This would allow to understand how the virus tolerate mutations at different sites and might provide information on the importance of different genomic regions for different stages of the infection.

## Acknowledgments

All authors wish to thank the www.gisaid.org and all the researchers that contributed their sequences to the database for sharing fundamental data for research. We think collaboration is the only approach to counteract the spread of SARS-CoV-2 and other similar endeavors. References for all sequences used in this work are in Supplementary Table 1. Authors from University of Milan wish to thank Fondazione Romeo ed Enrica Invernizzi for financial support. DS thanks the Italian Ministry of Education, University and Research (MIUR): for the Dipartimenti di Eccellenza Program (2018–2022) – Dept. of Biology and Biotechnology “L. Spallanzani”, University of Pavia.

## Notes

### Competing Interest Statement

The authors have declared no competing interest.

### Summary of Updates

Text has drastically changed as figures.

