## Supplementary figures and images for "Insurgence and worldwide diffusion of genomic variants in SARS-CoV-2 genomes"

### Supplemental Figure 1

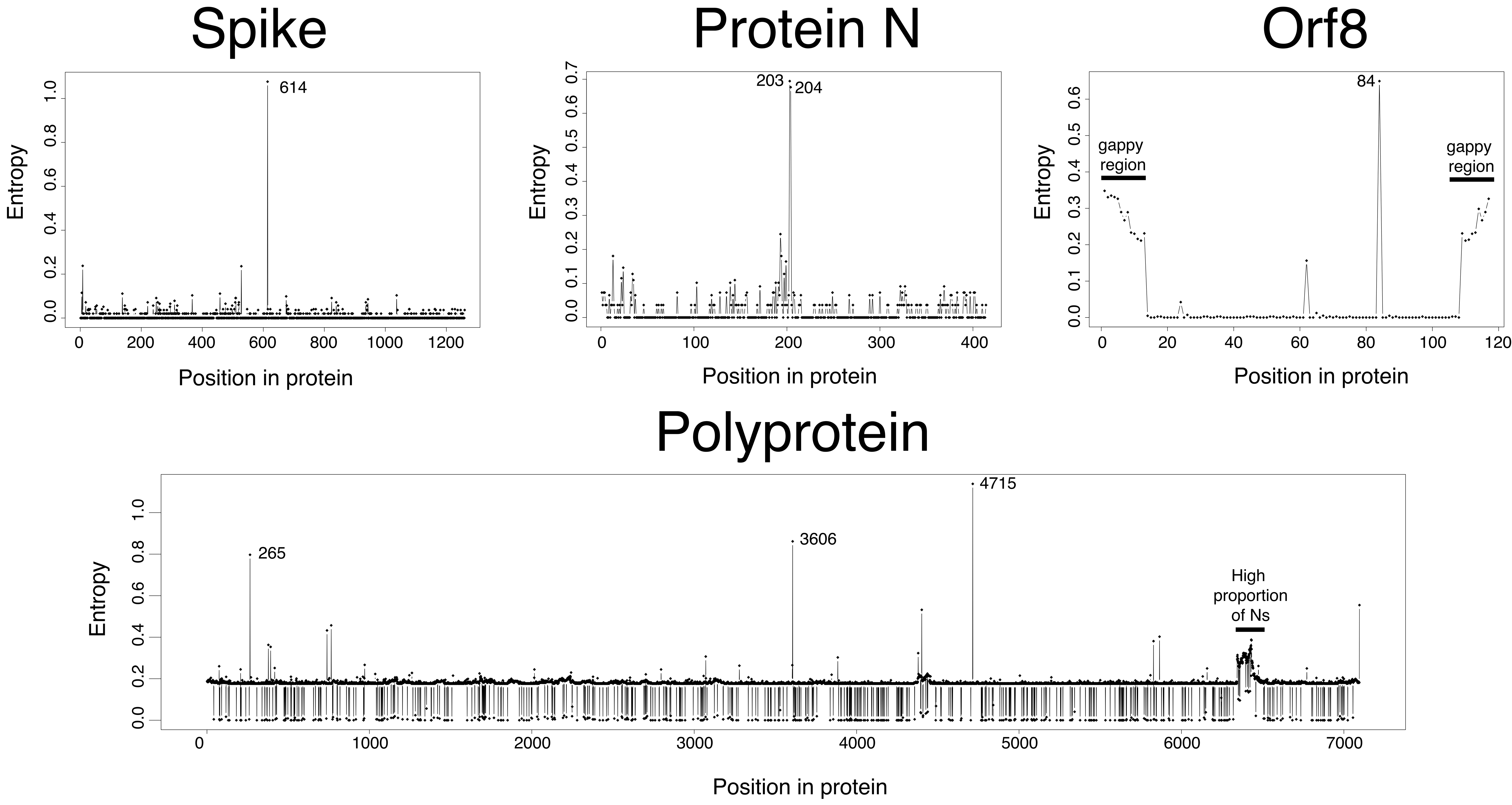

### Supplemental Figure 2

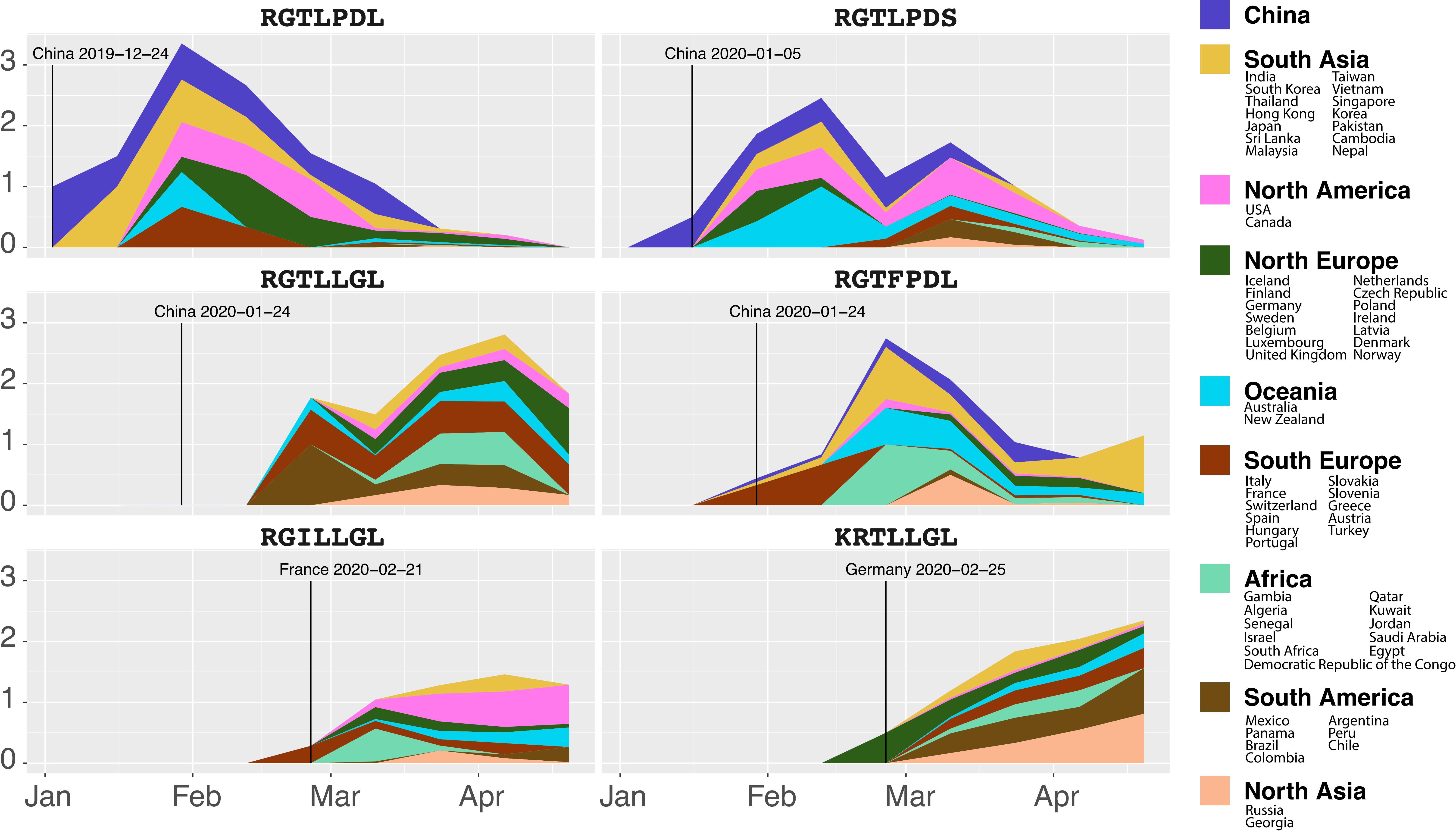
